# MARISMa: a routine MALDI-TOF MS dataset from 2018 to 2024 from Spain

**DOI:** 10.1101/2025.05.31.657186

**Authors:** Lucía Schmidt-Santiago, Inés López-Mareca, Mario Blázquez-Sánchez, José Miguel Moreno, Carlos Sevilla-Salcedo, Vanessa Gómez-Verdejo, Belén Rodríguez-Sánchez, David Rodríguez-Temporal, Alejandro Guerrero-López

**Affiliations:** Department of Signal and Communication Theory, Universidad Carlos III de Madrid, 28911 Spain; Department of Clinical Microbiology and Infectious Diseases, Gregorio Marañón Health Research Institute, Madrid, 28007 Spain; Computer Science and Engineering Department, Universidad Carlos III de Madrid, 28911 Spain

## Abstract

Clinical microbiology laboratories play a crucial role in identifying pathogens, guiding antibiotic treatment, and managing antimicrobial resistance (AMR). Matrix-Assisted Laser Desorption/Ionization Time-of-Flight Mass Spectrometry (MALDI-TOF MS) has become essential for rapid, accurate, and cost-effective microbial identification. Recent advances in integrating MALDI-TOF MS with Artificial Intelligence (AI) show promise in improving microbial detection and prediction of AMR. However, progress is limited by the lack of comprehensive and openly accessible datasets that restrict the validation, reproducibility, and applicability of the model.

To address this gap, we introduce a publicly available MALDI-TOF MS dataset comprising 202,700 unique spectra from isolates collected between 2018 and 2024 at the Hospital General Universitario Gregorio Marañón, Spain. This dataset includes 186,213 bacteria, 16,163 fungal, and 371 mycobacterial samples, of which 29,679 contain AMR annotations. This resource is openly and freely shared, rigorously curated, and designed to support a wide range of machine learning. By ensuring unrestricted access to high-quality, standardized data, this dataset aims to promote transparency, reproducibility, comparative benchmarking, and collaborative progress in AI-driven clinical microbiology.

## Background & Summary

The accurate and timely identification of microorganisms is crucial in clinical microbiology for effective patient management and infection control. Matrix-Assisted Laser Desorption/Ionization Time-of-Flight Mass Spectrometry (MALDI-TOF MS) has revolutionized this field by offering rapid, precise, and cost-effective identification of a broad spectrum of bacteria and fungi. This technology generates unique spectral fingerprints by ionizing proteins from microbial isolates. Its high reproducibility and efficiency have made MALDI-TOF MS indispensable in clinical diagnostics, proteomics, and biomarker discovery.

Despite these advances, significant limitations persist, particularly in regard to the identification of filamentous fungi and less common bacterial species [1, 2, 3]. Commercial datasets often lack comprehensive spectral data for these organisms, leading to challenges in accurate identification and potential misdiagnosis. For instance, the primary limitation in identifying filamentous fungi using MALDI-TOF MS is the limited reference spectra available, where the dataset of Resistance Information on Antimicrobials and MALDI-TOF Mass Spectra (DRIAMS) offers 5,203 fungi MALDI-TOF samples [4]. This gap underscores the need for more inclusive and representative spectral datasets to enhance the diagnostic capabilities of MALDI-TOF MS in clinical settings.

The integration of Artificial Intelligence (AI) techniques with MALDI-TOF MS data has significantly advanced microbial identification and characterization. Recent studies utilizing publicly accessible datasets, such as DRIAMS [4, 5], the University Medical Center Göttingen (MS-UMG) [6], and the Robert Koch Institute (RKI) [7] datasets, have demonstrated considerable improvements in microbial detection and antimicrobial resistance (AMR) prediction through the application of machine learning techniques [8, 9, 10, 11, 12, 13, 14]. For example, [8] we used DRIAMS along with a private dataset to train an MLP to predict AMR in *Pseudomonas aeruginosa*. Similarly, in [9] the authors combined the DRIAMS public *Staphylococcus aureus* dataset with a private dataset from Taiwan to train a LightGBM model to predict AMR. Other researchers, such as [10], propose using DRIAMS as a pretraining dataset to then finetune their own private dataset for AMR prediction. Despite these advancements, current publicly available datasets remain limited in number and scope, significantly hindering further developments in this rapidly growing research domain. In our systematic review [15] of 93 studies, we revealed that only 10% of them provide open access to their datasets, and only 5% of the total accompany these with publicly shared machine learning implementations or raw spectral files in the original Bruker Daltonics XMASS format, or *BrukerRaw* for short [16, 17].

While certain studies have shared datasets alongside their research for reproducibility, these datasets often lack comprehensive dataset papers or thorough metadata descriptions, reducing their practical utility for novel research initiatives. Furthermore, some publicly available data have already been preprocessed, thus constraining researchers solely to reproducing published findings and limiting their potential to explore novel hypotheses [18, 19, 20, 21, 22, 23, 24]. In contrast, numerous investigations rely on proprietary datasets or small datasets with limited representativeness [25, 26, 27, 28, 29, 30, 31], severely restricting reproducibility and generalizability.

Moreover, the shortage of openly accessible, well-characterized MALDI-TOF datasets critically impedes robust evaluation of AI models, particularly with respect to out-of-distribution (OOD) validation which is often lacked in the literature [13, 26, 25, 28, 24, 29, 22, 32, 33, 34]. Without open datasets, researchers are unable to rigorously test and demonstrate the transferability and clinical applicability of their models to external, heterogeneous data.

The open availability of both raw MALDI-TOF MS spectral data is essential for advancing reproducible research and fostering innovation in machine learning applications. This scenario underscores a persistent gap in the field: the lack of a large, standardized, and fully open-access databases that cover the full spectrum of microorganisms, including bacteria, fungi, and mycobacteria.

In light of these limitations, we present here the most extensive MALDI-TOF MS dataset to date: *MARISMa* (MALDI-TOF MS and Antimicrobial Resistance of Infectious Species at Marañón). This dataset comprises 202,700 unique spectra, which include data from 186,213 bacterial, 16,163 fungal, and 371 mycobacterial isolates. All spectra are provided in the original *BrukerRaw* format, ensuring that complete metadata regarding acquisition time, device information, and calibration details are available to researchers. This dataset is intended to support a wide range of research applications and to set a new benchmark for open-access resources in microbiology. Creating and openly sharing these resources will facilitate: (i) development of innovative and robust AI-driven methodologies, (ii) rigorous and comprehensive out-of-distribution validation studies, (iii) standardized comparative analysis across models and research groups, and ultimately, (iv) acceleration of collaborative advancements in AI-driven clinical microbiology research.

## Methods

A description of several aspects of the dataset is presented next, including the ethics declaration, a description of the data acquisition protocol, the sample protocol, the metadata acquisition, and antimicrobial resistance (AMR) availability.

### Ethics declaration

The study design and objectives were reviewed and approved by the Ethics Committee of the Gregorio Marañón Health Research Institute (CEIm). The Committee determined that the research met all applicable regulatory and ethical criteria for a waiver of informed consent, as it involved analysis of protein spectra derived from microbiological samples and did not involve direct use of human biological materials or identifiable human data.

Patients’ information was completely removed, and a unique identifier per bacteria was kept which is completely different from the one used in the Hospital’s electronic records. No personal data was exchanged with external researchers who had access to the dataset.

### Study selection criterion

This dataset includes all routine MALDI-TOF MS spectra acquired at Hospital General Universitario Gregorio Marañón between May 1, 2018 and December 31, 2024. The year 2018 was selected as the starting point to ensure consistency in acquisition instrumentation, as this marks the installation of the current MALDI-TOF device at the hospital.

Spectra were included if they originated from clinical routine workflows and were successfully assigned to a genus or species by the clinical microbiology laboratory. Exclusion criteria comprised (i) spectra flagged as low-quality (score below 1.7) by internal quality control systems or visual inspection, and (ii) spectra from unidentified or unclassified isolates.

To ensure the taxonomic reliability of species annotations assigned to each spectrum, a post-processing validation step was applied to the taxonomic identification outputs generated by the MALDI Biotyper system (Bruker Daltonics). Each processed sample yields both the raw spectrum in the *BrukerRaw* format and an associated `.xml` file containing the taxonomic results for each of the analytes, individual spectra acquired from different wells of the MALDI target plate.

Within each XML file, the Biotyper software returns a ranked list of up to 10 potential identifications per analyte, ordered by their `globalMatchValue`, a numerical score reflecting the similarity between the measured spectrum and entries in the reference library. Each result includes a genus and species-level classification, reported under the `referencePatternName` field.

The following filtering strategy was applied to retain only high-confidence and taxonomically consistent identifications:

1. **Confidence threshold check:** The first match (Rank 1) was evaluated. If its `globalMatchValue` was below a threshold of **1.7**, the analyte was **discarded**, as the classification was considered unreliable.
2. **Genus consistency check:** The genus of the second match (Rank 2) was compared with that of the first. If they differed, the analyte was **discarded**, indicating taxonomic inconsistency at the genus level.
3. **Species concordance:**
  - If the genus matched but the species differed, the `globalMatchValue` of the second match was evaluated:
    **–** If it was **greater than 1.7**, the analyte was **discarded**, due to conflicting high-confidence species-level identifications.
    **–** If it was **less than 1.7**, the analyte was **retained**, assuming the conflict was minor and not strongly supported.
4. **Third match validation:**
  - The genus must match that of the first two entries. If not, the analyte was **discarded**.
  - If the species differed:
    **–** A globalMatchValue above the threshold led to **rejection**.
    **–** A value below the threshold allowed the analyte to be **accepted**.

This multi-stage filtering process was systematically applied to all spectra in the dataset to maximize the accuracy and consistency of genus and species labels. By enforcing these criteria, we ensured that only spectra with coherent, high-confidence taxonomic annotations were retained in the final dataset.

It is important to note that each well-level spectrum (analyte) is individually annotated following this procedure. However, since multiple wells (analytes) correspond to the same original sample, discrepancies in their assigned genus or species can arise. In cases where wells from the same sample were classified into different taxonomic groups, the entire spectrum (i.e., the complete sample) was removed from the dataset to maintain label consistency and ensure data integrity.

An overview of this validation process is illustrated in Figure 1.

**Figure 1:**
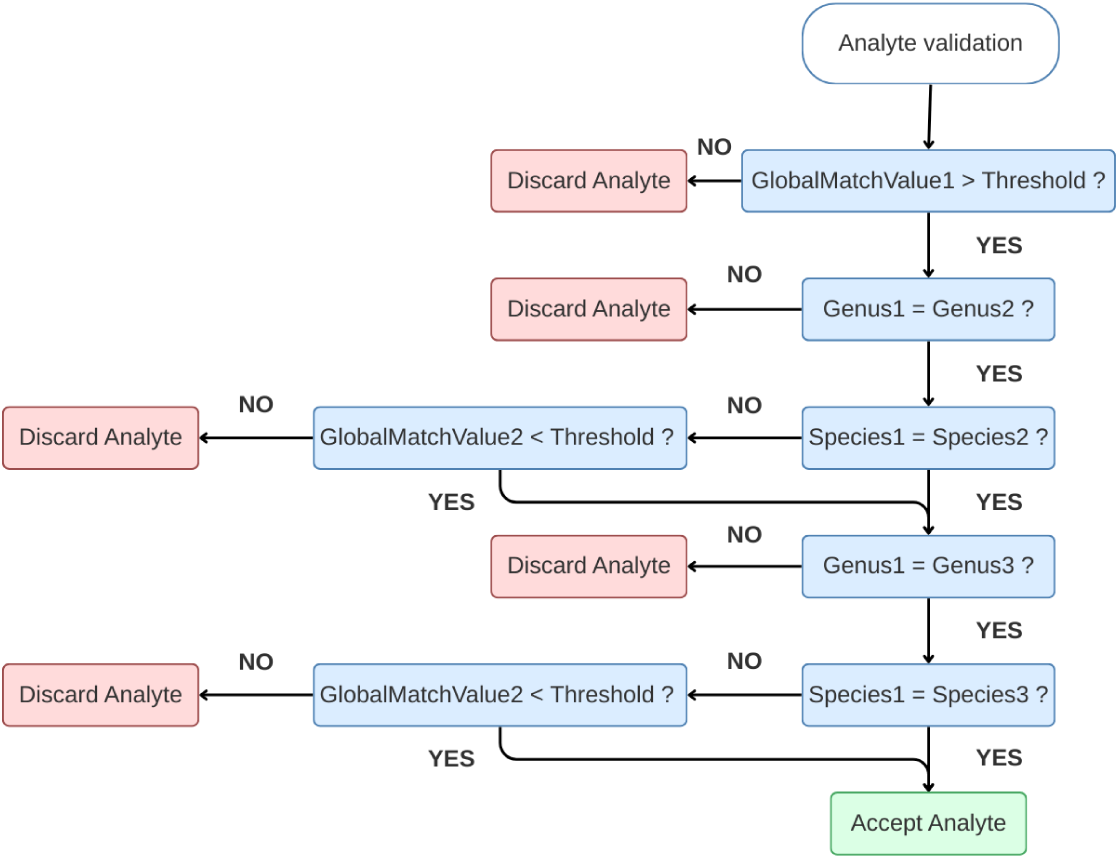
Flowchart describing the analyte validation procedure applied during the preprocessing of the *MARISMa* dataset. Only analytes with consistent genus and species predictions among the top-3 classification matches, and with high-confidence global match values (threshold *≥* 1.7), are retained.

The final dataset comprises 186,213 bacterial, 16,163 fungal, and 371 mycobacterial isolates, all shared in their raw spectral form. Each isolate may have multiple spectral replicates, generated as part of routine reproducibility testing; in total, the dataset includes 241,980 MALDI-TOF spectra. Figure 2a highlights the most represented microorganisms in the collection, which predominantly belong to bacterial species classified as WHO critical or high-priority pathogens [35]. These include *Escherichia coli* and *Klebsiella pneumoniae* (critical priority), as well as *Enterococcus faecalis* and *Staphylococcus aureus* (high priority).

**Figure 2:**
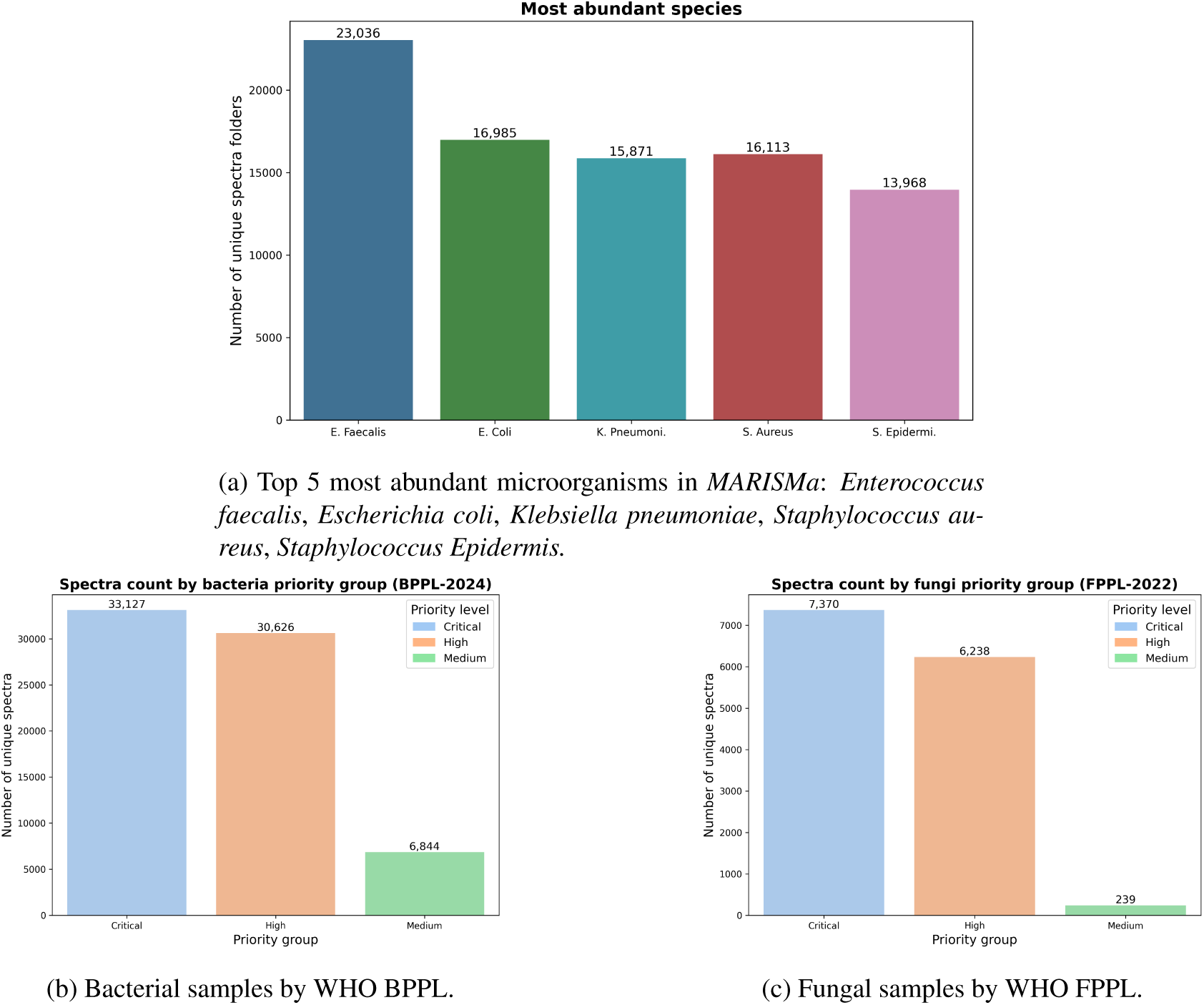
Overview of sample distribution in *MARISMa*.

For bacterial species, we reference the 2024 WHO Bacterial Priority Pathogens List (BPPL) [35], which categorizes pathogens into three priority tiers—critical, high, and medium—based on their public health relevance and the urgency for research and antimicrobial development. Our dataset thus contains representatives from all bacterial families listed in the WHO classification, as detailed in Figure 2b. Although we do not provide antibiotic susceptibility profiles for all samples, only for 29, 679 samples in total, we include isolates identified as belonging to these species regardless of resistance phenotype.

For the fungal species, we follow the WHO Fungal Priority Pathogens List (FPPL) published in 2022 [36], which identifies 19 fungal pathogens of concern and similarly groups them into critical, high, and medium priority categories. Our dataset includes representative isolates from all fungal families listed in this classification as detailed in Figure 2c.

By encompassing all WHO-designated bacterial and fungal priority pathogens, this dataset offers an exceptionally valuable resource for the development, benchmarking, and evaluation of AI-driven tools in clinical microbiology, AMR surveillance, and pathogen detection.

### Sample protocol

Protein spectra included in the *MARISMa* dataset were acquired from isolates sent for routine identification to the Clinical Microbiology and Infectious Diseases department of the Hospital General Universitario Gregorio Marañón. One spectrum was acquired per microbial isolate using a mass spectrometer MBT Smart MALDI Biotyper (Bruker Daltonics, Bremen, Germany). The instrument was calibrated daily using a commercial bacterial test standard (Bruker Daltonics), prior to spectra acquisition. Isolates were cultured on standard media for bacterial and fungal isolates. The spectra from microorganisms analyzed directly from blood cultures were obtained using a commercial kit (Sepsityper kit, Bruker Daltonics) that concentrates the microorganisms present in 1 ml of blood culture and removes blood components using a washing buffer.

### Metadata

*BrukerRaw* spectra are composed of several files, mainly `fid`, `acqu` and `proc`. The first is a binary file with the intensity values of the spectrogram, and the last two are plain text files containing metadata that characterizes the acquired sample and the instrument used. Among the hundreds of variables found between this pair of files alone, we highlight a dozen that are useful to researchers:

- **Sample ID (`$ID_raw`).** Random, unique identifier automatically generated during acquisition.
- **Target ID (`$TgIDS`).** Same as sample ID, but identifies the whole group of samples acquired at once in the same batch.
- **Sample position (`$PATCHNO`).** Acquired position inside the plate (e.g., A1).
- **Acquisition date (`$AQ_DATE`).** Timestamp for when the sample was acquired.
- **Calibration date (`$CLDATE`).** Timestamp of the last calibration of the acquisition instrument.
- **Instrument serial number (`$InstrID`).** Unique identifier of the acquisition instrument set by the manufacturer.
- **Instrument type (`$InstTyp`).** Numeric code determining the type of instrument (e.g., type 9 is microflex). As with most of the *BrukerRaw* format, these constants are not officially documented, but the research community has already taken on this task [37].
- **Digitizer type (`$DIGTYP`).** Same as instrument type, but for the digitizer (e.g., type 19 is Bruker BD0G5).
- **flexControl version (`$FCVer`).** Acquisition software version.
- **AIDA version (`$Acquver`).** Calibration software version.
- **Spectrum size (`$TD`).** Number of spectrum peaks (intensity time periods).
- **Spectrum delay (`$DELAY`).** Time in nanoseconds until the first measured intensity.
- **Sample rate (`$DW`).** Nanoseconds between each measured intensity.
- **Mass calibration constants (`$ML1`, `$ML2`, `$ML3`).** Also referred to as dwell times. Parameters needed to convert measured intensities from the `fid` file to pairs of m/z using the formula described in [38].

## Data Record

Data access is facilitated through both the Zenodo repository [39] version 2.0.0. The structural representation of the dataset is illustrated in Fig. 3 and Fig. 4. The dataset comprises 202,700 unique MALDI-TOF MS spectra acquired from bacterial and fungal isolates, all provided in the original *BrukerRaw* format.

**Figure 3:**
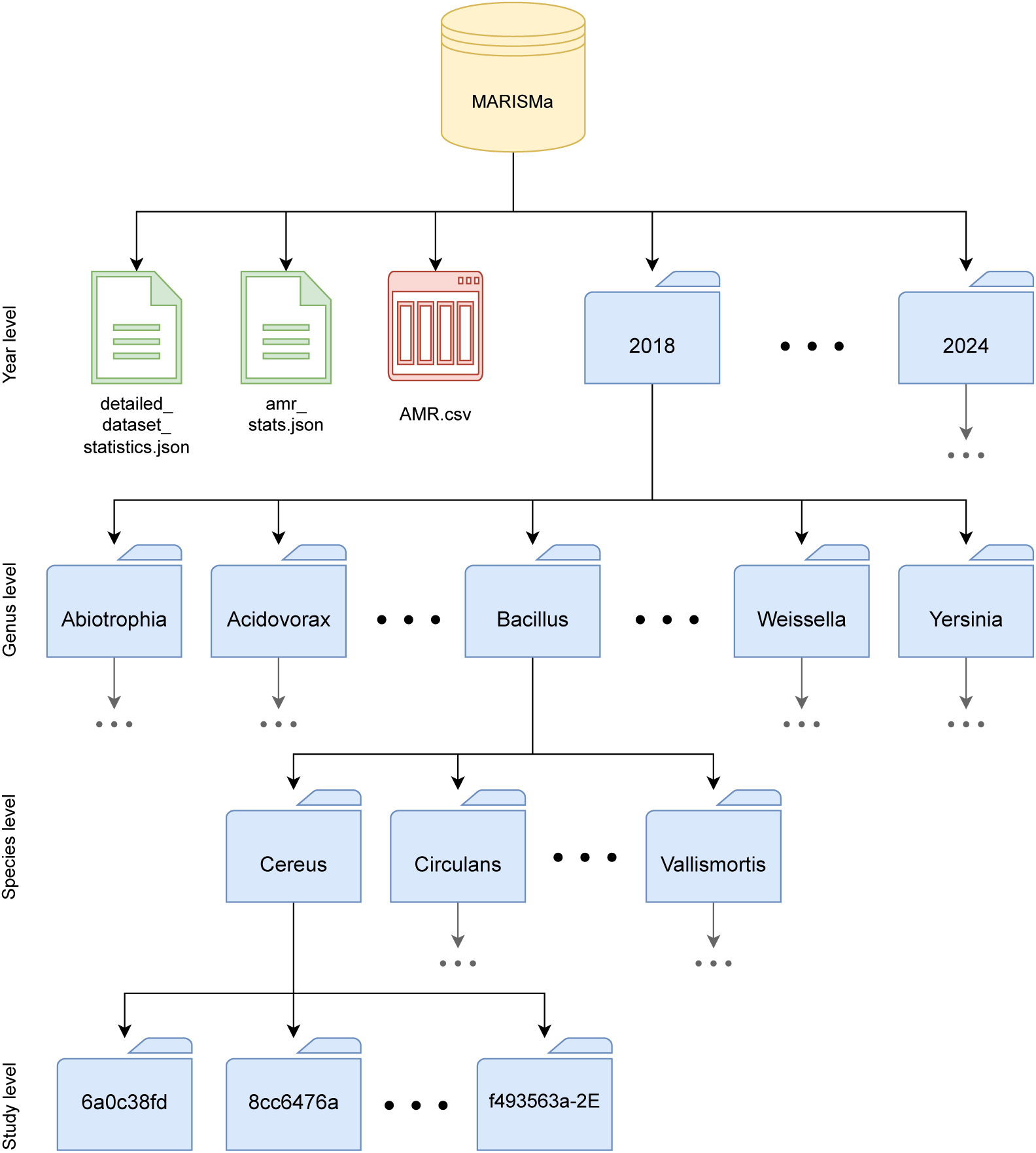
Hierarchical folder structure of the *MARISMa* dataset. The dataset is organized chronologically, with one folder per acquisition year (`2018` to `2024`). Within each year, data is grouped by bacterial genus, then by species, and finally by study. At the root level of the dataset, two files — `detailed_dataset_statistics.json` and `amr_stats.json` — provide global summary statistics: the former reports the distribution of spectra per genus and species, while the latter summarizes antimicrobial resistance (AMR) labels by antibiotic and species. Additionally, AMR labels are provided for 29,679 isolates in `AMR.csv`.

**Figure 4:**
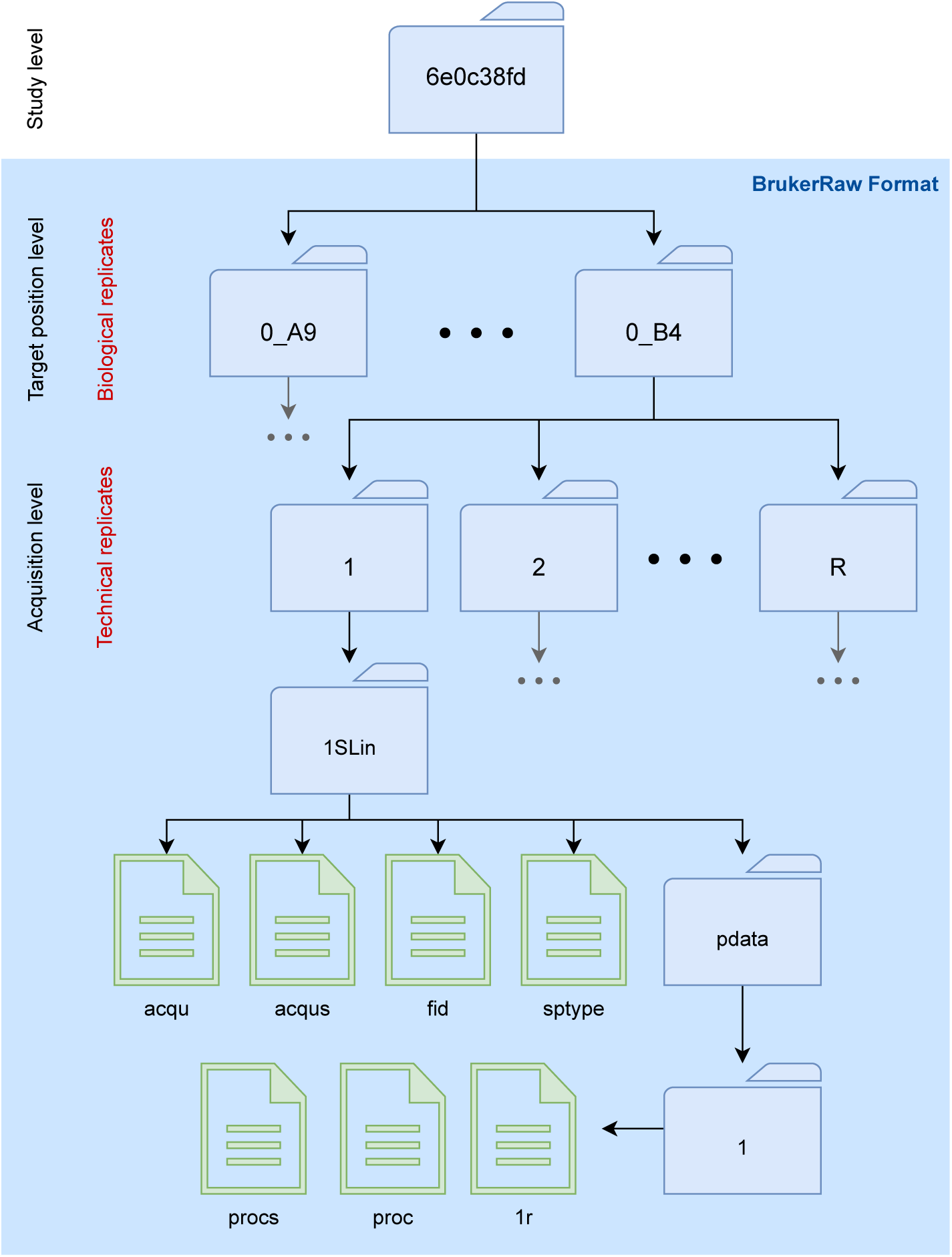
Structure of the raw data format used in the *MARISMa* dataset, following the BrukerRaw standard. Study folders contain subfolders representing MALDI target plate positions (e.g., `0_A1`, `0_B8`), which correspond to biological replicates. Each target position folder may include multiple acquisition folders (`1`, `2`, . . . ) representing technical replicates. These replicate folders contain raw spectra in Bruker format (within a `1SLin` structure) with raw acquisition files and a `pdata` folder. Relevant files are the `fid` file, which stores the mass spectrum in binary format, and the `acqu` file, which contains essential metadata such as the acquisition and calibration dates. Other files like `acqus` and `sptype` contain complementary acquisition settings. This structure ensures reproducibility and traceability of each spectral acquisition.

This format ensures that complete metadata—such as acquisition time, device information, and calibration details—is available to researchers, thereby facilitating robust reproducibility and further data-driven investigations.

The dataset is systematically organized to facilitate efficient data retrieval and analysis, firstly divided by year of data acquisition (2018–2024). Within each year, data is grouped by bacterial genera, and within each genus, by species. For each species, the dataset contains the studies in which that species was analyzed, identified by an anonymized identifier. Within each study folder, data is organized following *the BrukerRaw* format, i.e., subfolders named according to the target position on the MALDI target plate (e.g., `0_A1`, `0_A2`, `0_D5`), which correspond to biological replicates. Within each target position folder, there is always a folder named `1`, representing the first acquisition of the sample; when experiments are repeated on the same sample, additional folders named `2`, `3`, etc., appear, corresponding to technical replicates. Each acquisition folder contains the standard *BrukerRaw* structure: a `1SLin` folder with acquisition files (`acqu`, `acqus`, `fid`, `sptype`) and a `pddata` folder with additional metadata. AMR labels are provided in a dedicated `csv` file. Statistics files summarize the number of genera, species, studies, spectra, and antibiotics available.

**Table 1:**
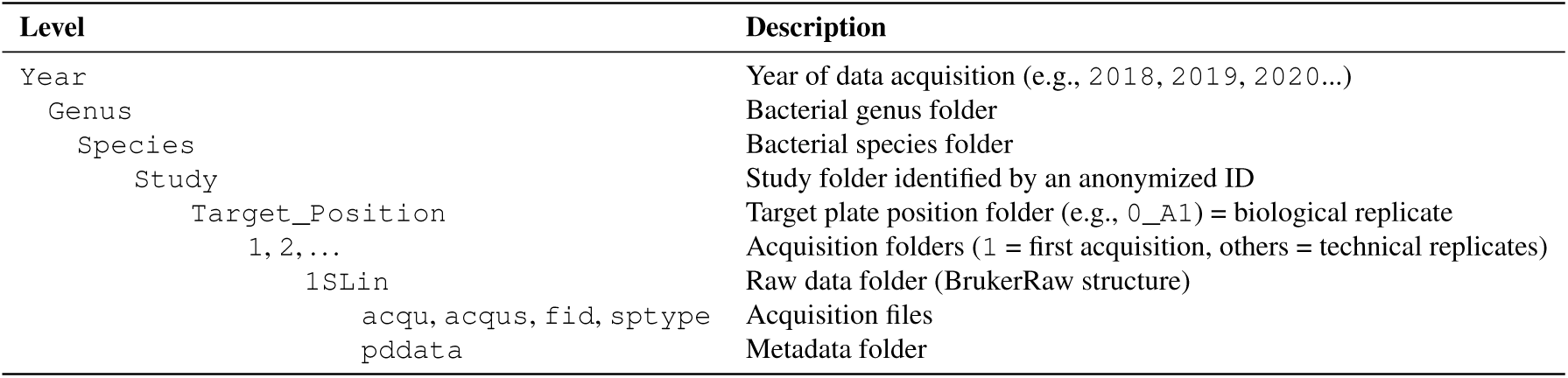
Folder structure of the *MARISMa* dataset.

### AMR labels

We provide AMR annotations for 29, 679 unique isolates, including susceptibility results for up to 78 antibiotics for 220 different species. Table 2 shows the AMR analysis, showing counts of resistant (R), susceptible (S), and intermediate (I) phenotypes based on routine clinical testing.

**Table 2:** Summary of available AMR labels per antibiotic in MARISMa. ’Species’ indicates the number of species with available AMR labels for the given antibiotic. Resistant (R), Susceptible (S), and Intermediate (I) columns show absolute counts and percentages.

| Antibiotic | Species | R (%) | S (%) | I (%) | Total |
| --- | --- | --- | --- | --- | --- |
| Amikacin | 115 | 5121 (18%) | 23794 (82%) | 1 (0%) | 28916 |
| Amoxicillin | 5 | 4 (27%) | 11 (73%) | 0 (0%) | 15 |
| Amoxicillin/Clavulanic acid | 119 | 9126 (35%) | 17227 (65%) | 3 (0%) | 26356 |
| Amphotericin B | 6 | 2 (5%) | 39 (95%) | 0 (0%) | 41 |
| Ampicillin | 145 | 26440 (79%) | 6941 (21%) | 13 (0%) | 33394 |
| Ampicillin/Sulbactam | 78 | 9239 (51%) | 8335 (46%) | 674 (4%) | 18248 |
| Anidulafungin | 8 | 0 (0%) | 99 (99%) | 1 (1%) | 100 |
| Azithromycin | 18 | 109 (20%) | 435 (80%) | 0 (0%) | 544 |
| Aztreonam | 84 | 5335 (23%) | 16098 (71%) | 1333 (6%) | 22766 |
| Caspofungin | 8 | 1 (1%) | 91 (94%) | 5 (5%) | 97 |
| Cefalotin | 1 | 593 (29%) | 1473 (71%) | 0 (0%) | 2066 |
| Cefazolin | 8 | 1247 (90%) | 145 (10%) | 0 (0%) | 1392 |
| Cefepime | 97 | 5841 (24%) | 17051 (70%) | 1387 (6%) | 24279 |
| Cefiderocol | 6 | 28 (33%) | 55 (64%) | 3 (3%) | 86 |
| Cefixime | 51 | 4686 (25%) | 13722 (75%) | 0 (0%) | 18408 |
| Cefotaxime | 136 | 4664 (21%) | 17531 (79%) | 56 (0%) | 22251 |
| Cefoxitin | 43 | 2860 (47%) | 3113 (52%) | 63 (1%) | 6036 |
| Ceftaroline | 8 | 20 (1%) | 2004 (99%) | 0 (0%) | 2024 |
| Ceftazidime | 84 | 3345 (16%) | 16199 (77%) | 1621 (8%) | 21165 |
| Ceftazidime/Avibactam | 71 | 434 (2%) | 19898 (98%) | 0 (0%) | 20332 |
| Ceftolozane/Tazobactam | 68 | 2063 (10%) | 18239 (90%) | 0 (0%) | 20302 |
| Ceftriaxone | 14 | 11 (4%) | 299 (96%) | 2 (1%) | 312 |
| Cefuroxime | 66 | 9677 (42%) | 10456 (46%) | 2733 (12%) | 22866 |
| Cephalexin | 56 | 4798 (42%) | 6631 (58%) | 0 (0%) | 11429 |
| Chloramphenicol | 103 | 2401 (23%) | 7873 (77%) | 11 (0%) | 10285 |
| Ciprofloxacin | 181 | 10018 (31%) | 18429 (57%) | 3692 (11%) | 32139 |
| Clarithromycin | 20 | 108 (33%) | 220 (67%) | 0 (0%) | 328 |
| Clindamycin | 98 | 4676 (66%) | 2374 (34%) | 0 (0%) | 7050 |
| Clotrimazole | 2 | 1 (25%) | 3 (75%) | 0 (0%) | 4 |
| Colistin | 82 | 2602 (11%) | 21052 (89%) | 0 (0%) | 23654 |
| Dalbavancin | 3 | 10 (62%) | 6 (38%) | 0 (0%) | 16 |
| Daptomycin | 42 | 70 (1%) | 8425 (99%) | 0 (0%) | 8495 |
| Doripenem | 7 | 0 (0%) | 109 (100%) | 0 (0%) | 109 |
| Doxycycline | 38 | 87 (17%) | 433 (83%) | 0 (0%) | 520 |
| Ertapenem | 85 | 3832 (16%) | 20208 (84%) | 5 (0%) | 24045 |
| Erythromycin | 85 | 6380 (70%) | 2752 (30%) | 0 (0%) | 9132 |
| Ethambutol | 1 | 0 (0%) | 2 (100%) | 0 (0%) | 2 |
| Fluconazole | 7 | 11 (11%) | 47 (48%) | 39 (40%) | 97 |

| Antibiotic | Species | R (%) | S (%) | I (%) | Total |
| --- | --- | --- | --- | --- | --- |
| Fosfomycin | 140 | 7782 (25%) | 23488 (75%) | 0 (0%) | 31270 |
| Fusidic acid | 14 | 534 (45%) | 656 (55%) | 0 (0%) | 1190 |
| Gentamicin | 111 | 8520 (28%) | 21584 (72%) | 0 (0%) | 30104 |
| Gentamicin 500 | 14 | 1384 (33%) | 2840 (67%) | 0 (0%) | 4224 |
| Imipenem | 127 | 1987 (7%) | 20108 (67%) | 8122 (27%) | 30217 |
| Isavuconazole | 4 | 2 (20%) | 8 (80%) | 0 (0%) | 10 |
| Isoniazid | 1 | 0 (0%) | 2 (100%) | 0 (0%) | 2 |
| Itraconazole | 3 | 0 (0%) | 9 (100%) | 0 (0%) | 9 |
| Levofloxacin | 164 | 9169 (28%) | 19441 (59%) | 4458 (13%) | 33068 |
| Linezolid | 73 | 81 (1%) | 8831 (99%) | 0 (0%) | 8912 |
| Mecillinam | 15 | 1 (0%) | 1493 (100%) | 0 (0%) | 1494 |
| Meropenem | 98 | 831 (3%) | 22750 (95%) | 270 (1%) | 23851 |
| Meropenem/Vaborbactam | 1 | 0 (0%) | 2 (100%) | 0 (0%) | 2 |
| Metronidazole | 8 | 0 (0%) | 19 (100%) | 0 (0%) | 19 |
| Micafungin | 8 | 0 (0%) | 97 (100%) | 0 (0%) | 97 |
| Minocycline | 47 | 3163 (29%) | 7650 (71%) | 35 (0%) | 10848 |
| Moxifloxacin | 32 | 901 (75%) | 296 (25%) | 2 (0%) | 1199 |
| Mupirocin | 23 | 525 (15%) | 2965 (85%) | 0 (0%) | 3490 |
| Nalidixic acid | 23 | 495 (74%) | 174 (26%) | 0 (0%) | 669 |
| Nitrofurantoin | 44 | 5161 (28%) | 12967 (72%) | 0 (0%) | 18128 |
| Norfloxacin | 58 | 9513 (50%) | 9599 (50%) | 7 (0%) | 19119 |
| Oritavancin | 2 | 14 (67%) | 7 (33%) | 0 (0%) | 21 |
| Oxacillin | 27 | 1197 (33%) | 2432 (67%) | 0 (0%) | 3629 |
| Penicillin | 101 | 3407 (36%) | 5734 (61%) | 198 (2%) | 9339 |
| Piperacillin | 63 | 18602 (88%) | 2339 (11%) | 267 (1%) | 21208 |
| Piperacillin/Tazobactam | 89 | 6382 (27%) | 16251 (68%) | 1325 (6%) | 23958 |
| Posaconazole | 6 | 0 (0%) | 42 (100%) | 0 (0%) | 42 |
| Pristinamycin | 46 | 2996 (34%) | 5697 (65%) | 64 (1%) | 8757 |
| Pyrazinamide | 1 | 0 (0%) | 2 (100%) | 0 (0%) | 2 |
| Rifampin | 47 | 3144 (71%) | 1297 (29%) | 1 (0%) | 4442 |
| Streptomycin | 3 | 2 (50%) | 2 (50%) | 0 (0%) | 4 |
| Tedizolid | 1 | 2 (100%) | 0 (0%) | 0 (0%) | 2 |
| Teicoplanin | 36 | 207 (3%) | 7043 (97%) | 0 (0%) | 7250 |
| Tetracycline | 48 | 4410 (50%) | 4316 (49%) | 15 (0%) | 8741 |
| Ticarcillin | 78 | 17301 (85%) | 2143 (11%) | 965 (5%) | 20409 |
| Tigecycline | 54 | 3371 (46%) | 3930 (54%) | 0 (0%) | 7301 |
| Tobramycin | 111 | 10349 (33%) | 21117 (67%) | 1 (0%) | 31467 |
| Trimethoprim | 61 | 3583 (54%) | 2985 (45%) | 108 (2%) | 6676 |
| Trimethoprim/Sulfamethoxazole | 169 | 10674 (35%) | 19086 (63%) | 437 (1%) | 30197 |

| Antibiotic | Species | R (%) | S (%) | I (%) | Total |
| --- | --- | --- | --- | --- | --- |
| Vancomycin | 102 | 251 (3%) | 9102 (97%) | 0 (0%) | 9353 |
| Voriconazole | 9 | 0 (0%) | 73 (99%) | 1 (1%) | 74 |

The most commonly tested antibiotics include beta-lactams, carbapenems, and fluoroquinolones. As expected, certain antibiotics, such as ampicillin, show nearly universal resistance, whereas others, such as colistin and imipenem, retain high susceptibility rates. These data illustrate the clinical relevance and variability present in the dataset and underscore its utility for AMR prediction tasks.

AMR data is provided in a `CSV` file, where each row corresponds to a spectrum, identified by an identifier in the first column. This identifier refers to an individual well on the MALDI target plate, so there may be different wells for the same isolate—meaning that all spectra (wells) derived from the same sample might not share the same AMR profile. In the folder structure, this identifier corresponds to a directory located under each `Species/` folder, containing all associated target well subfolders.

The `CSV` includes several metadata fields: sample identifier, target position (well), year of collection, full microorganism name, unified species name, anatomical source of the sample, and details on individual resistance mechanisms. AMR results follow in column pairs: each pair consists of the Minimum Inhibitory Concentration (MIC) for a given antibiotic and its corresponding clinical interpretation (e.g., R, S, I, and variants of these). Resistance profiles are provided for up to 78 antibiotics.

While the dataset includes AMR labels for multiple species, the distribution is highly unbalanced: out of 29,679 AMR-labeled samples, 14,201 correspond to *Klebsiella pneumoniae*. This imbalance is due to the comprehensive retrieval of these isolates across all available years (2018–2024), as part of an earlier clinical study. For other species, AMR data is currently limited to the year 2024, following a change in the hospital’s information system that enabled broader extraction of AST results.

The distribution of interpretation labels, R and S, across antibiotics is highly unbalanced and varies considerably between antibiotics and species (Table 2). Each antibiotic exhibits a distinct resistance profile across species, shaped by the natural prevalence of resistance mechanisms in clinical environments. This imbalance is not only a result of usage frequency but rather reflects biological and evolutionary factors — for some antibiotics, resistance genes are widespread, while for others, resistance mechanisms are rarely observed. Furthermore, certain antibiotics in *MARISMa* have extensive data spanning many species, while others include only a small number of samples, underscoring the presence of substantial missing data. This uneven distribution mirrors real-world clinical practices, where some antimicrobials are routinely tested and others are used only in specific cases. Figure 5 exemplifies this phenomenon by illustrating the distribution of resistance and susceptibility patterns for *Klebsiella pneumoniae* isolates, further highlighting the variability and imbalance inherent to the dataset.

**Figure 5:**
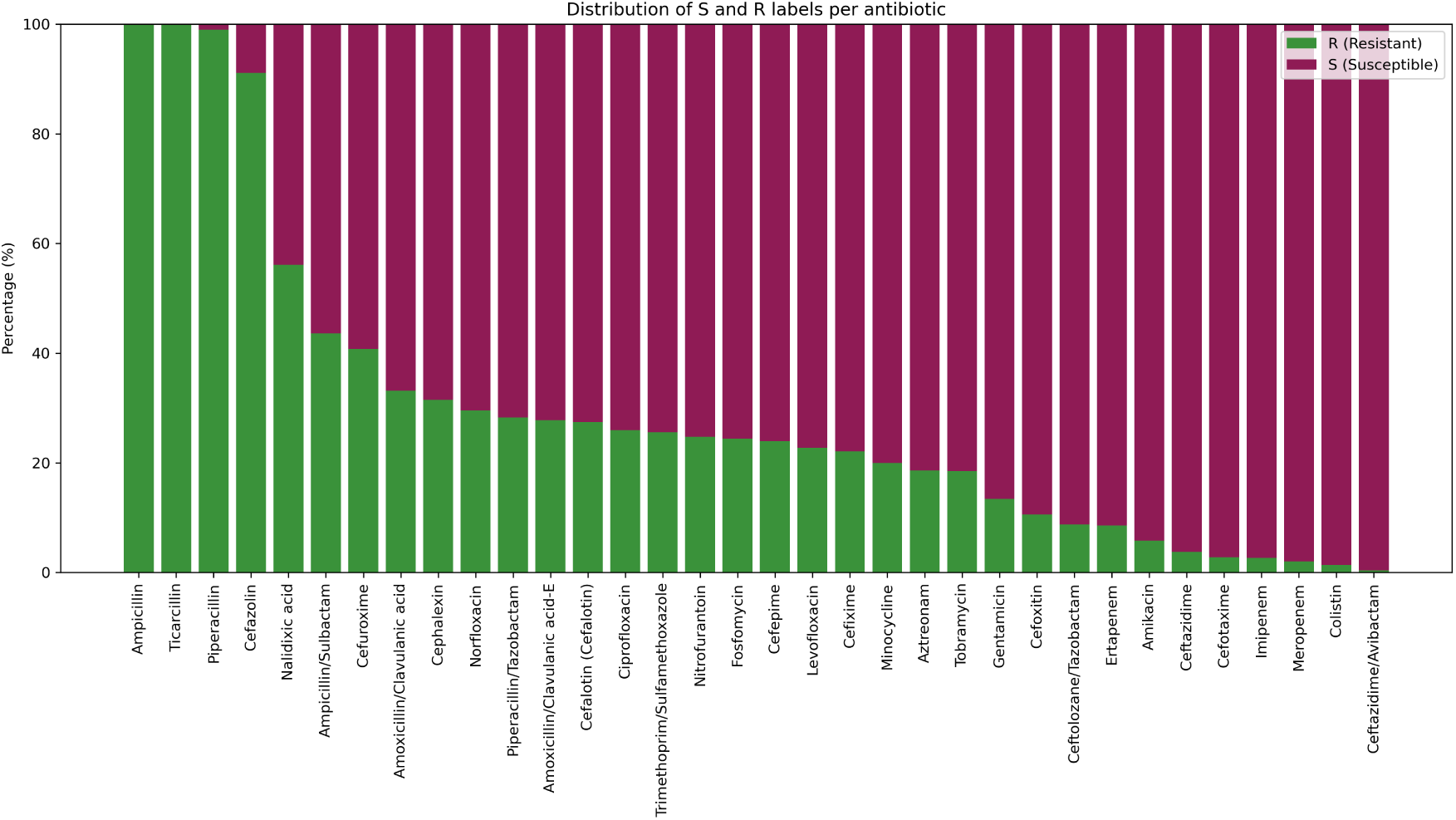
Distribution of resistant (R) and susceptible (S) labels across 34 different antibiotics for *Klebsiella pneumoniae* isolates, highlighting variability in resistance profiles. Antibiotics are ordered by decreasing resistance prevalence.

## Technical Validation

The *MARISMa* dataset has been partially examined in several studies, establishing its usefulness for different applications, such as antibiotic resistance prediction in *Klebsiella pneumoniae* [40], automatic ribotyping *of Clostridium difficile* [41], the development of data augmentation techniques for microbiology [42], and automatic surveillance of *Escherichia coli*[43], among others.

### Demonstration of dataset utility and reproducibility

To facilitate immediate use and reproducibility, we provide a demonstration notebook, `simple_classifier`, available in the associated GitHub repository. This notebook exemplifies how to query, preprocess, and classify spectra using our MALDI-TOF MS dataset.

The provided code includes a flexible function that allows users to construct training datasets by selecting specific acquisition years, genera, and species. For instance, by specifying years `[2018, 2019]` and genus `Klebsiella`, the function retrieves all matching samples and assembles a corresponding dataset, which can be used for downstream machine learning tasks.

Each spectrum is then preprocessed using standard signal processing techniques widely adopted in MALDI-TOF analysis, whose hyperparameters can be tuned by the user. For instance, the example given follows:

~~~
bin_step = 9
spec = SpectrumObject.from_bruker(acqu_file, fid_file)
spec = VarStabilizer(method=’sqrt’)(spec)
spec = Smoother(halfwindow=10, polyorder=3)(spec)
spec = BaselineCorrecter(method=’SNIP’, snip_n_iter=10)(spec)
spec = Normalizer()(spec)
spec = Trimmer(min=2000, max=20000)(spec)
spec = Binner(step=bin_step)(spec)
~~~

A simple `Random Forest` classifier is then trained to discriminate between all available *Klebsiella* species within the selected time frame. This example showcases not only the high quality and diversity of the dataset, but also its ease of use and adaptability. Users are encouraged to modify the notebook to select different time periods, genera, or species, allowing rapid prototyping and testing of new machine learning approaches.

#### Impact of preprocessing on spectral clustering

To evaluate the structure and interpretability of the MALDI-TOF MS spectra, we performed a t-SNE analysis of the top ten most represented organisms in the dataset, including bacterial and fungal species. Figure 6 illustrates the resulting embeddings before and after applying standard preprocessing steps.

**Figure 6:**
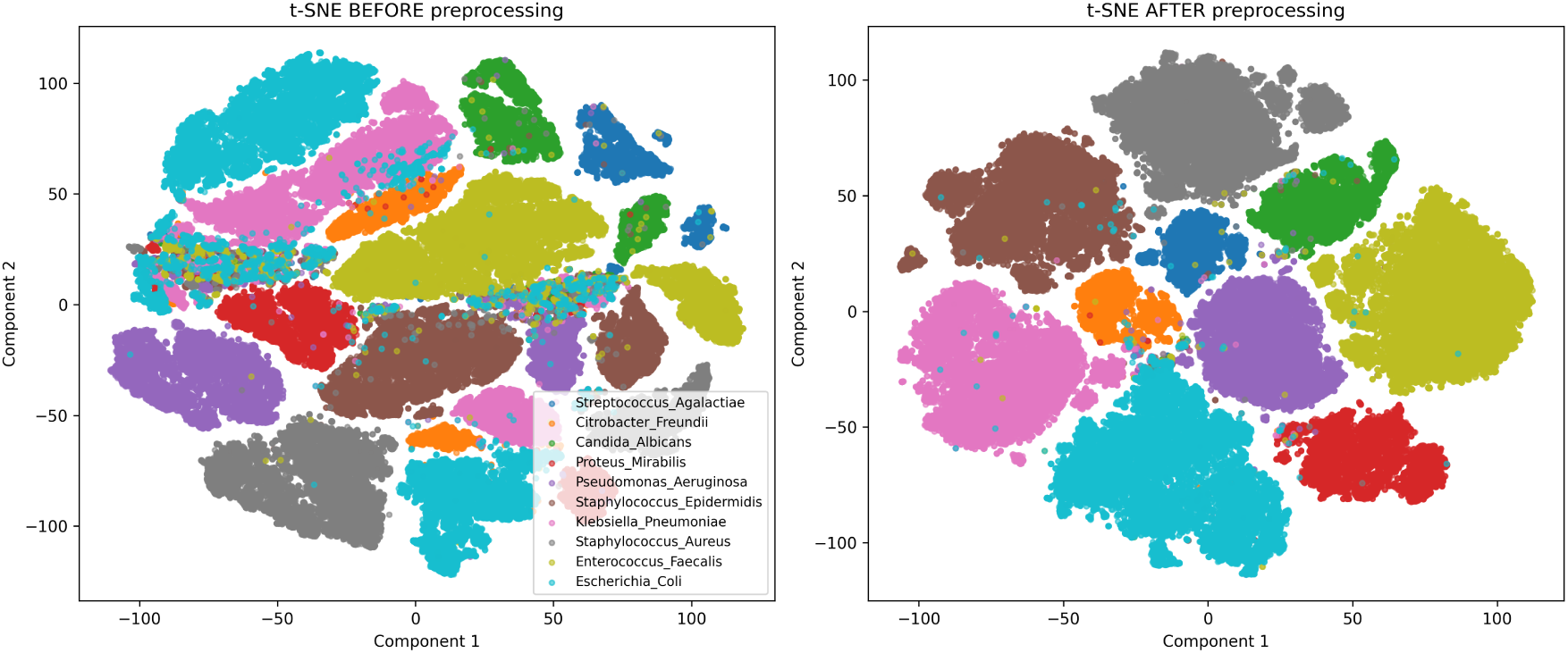
t-SNE visualizations of the top ten most represented microorganisms in the dataset. Left: Raw spectra as acquired, interpolated to equal length. Some overlap and misclassification are visible. Right: The same data after preprocessing, including variance stabilization, smoothing, baseline correction, normalization, and trimming. Well-separated clusters indicate improved inter-class structure and highlight the utility of preprocessing.

In the left panel, the t-SNE was computed on the raw spectra (interpolated to a common length) as they were originally acquired. While some clustering is evident, several species exhibit overlapping or poorly defined boundaries, with certain samples scattered across clusters. This highlights the challenges of working directly with unprocessed spectra, which may contain high variance, noise, or baseline drift.

In the right panel, we show the same dataset after applying a commonly adopted preprocessing pipeline in MALDI-TOF analysis. Specifically, each spectrum underwent variance stabilization, smoothing, baseline correction, normalization, and trimming. The resulting t-SNE reveals clearly defined clusters with minimal overlap, reflecting a more coherent biological structure in the data.

Although we release all spectra in raw form to ensure full flexibility and transparency, this comparison underscores the importance of preprocessing in downstream machine learning tasks and supports the use of established preprocessing pipelines for spectral feature extraction and classification.

#### Temporal consistency of spectra acquisition

To assess the robustness and temporal consistency of the MALDI-TOF MS data, we conducted a t-SNE 2D analysis for four of the most frequent bacterial species in the dataset. For each species, we visualized the t-SNE embedding of their spectra, color-coded by the year of acquisition (2018–2024).

#### Preservation of taxonomic separability across years

To further evaluate the diversity and consistency of the MALDI-TOF MS data across the study period, we performed an annual t-SNE analysis from 2018 to 2024. For each year, we extracted spectra corresponding to the ten most frequently represented microorganisms (including both bacteria and fungi) and applied dimensionality reduction to visualize their latent structure.

Figure 8 displays the resulting t-SNE plots for some year. Although each embedding was computed independently and is thus not directly comparable in position or scale, each year consistently yields ten wellseparated clusters. This repeated clustering pattern across years strongly supports the taxonomic distinguishability of the major organisms in the dataset and reflects both the biological diversity of the samples and the stable acquisition quality over time.

**Figure 7:**
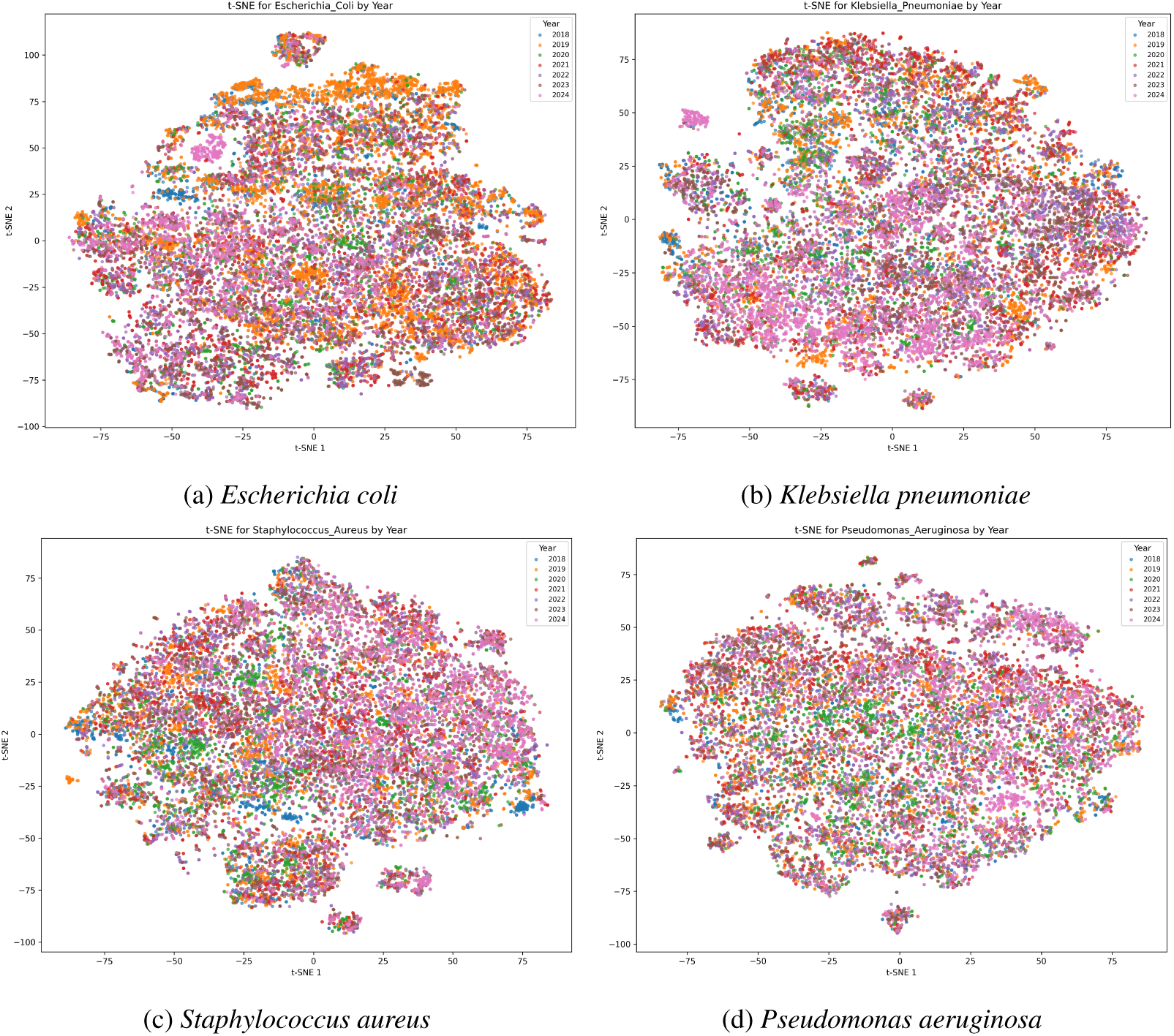
t-SNE visualization of MALDI-TOF spectra for four key bacterial species, color-coded by year of acquisition. The absence of temporal clustering indicates consistency in spectral quality across the years.

**Figure 8:**
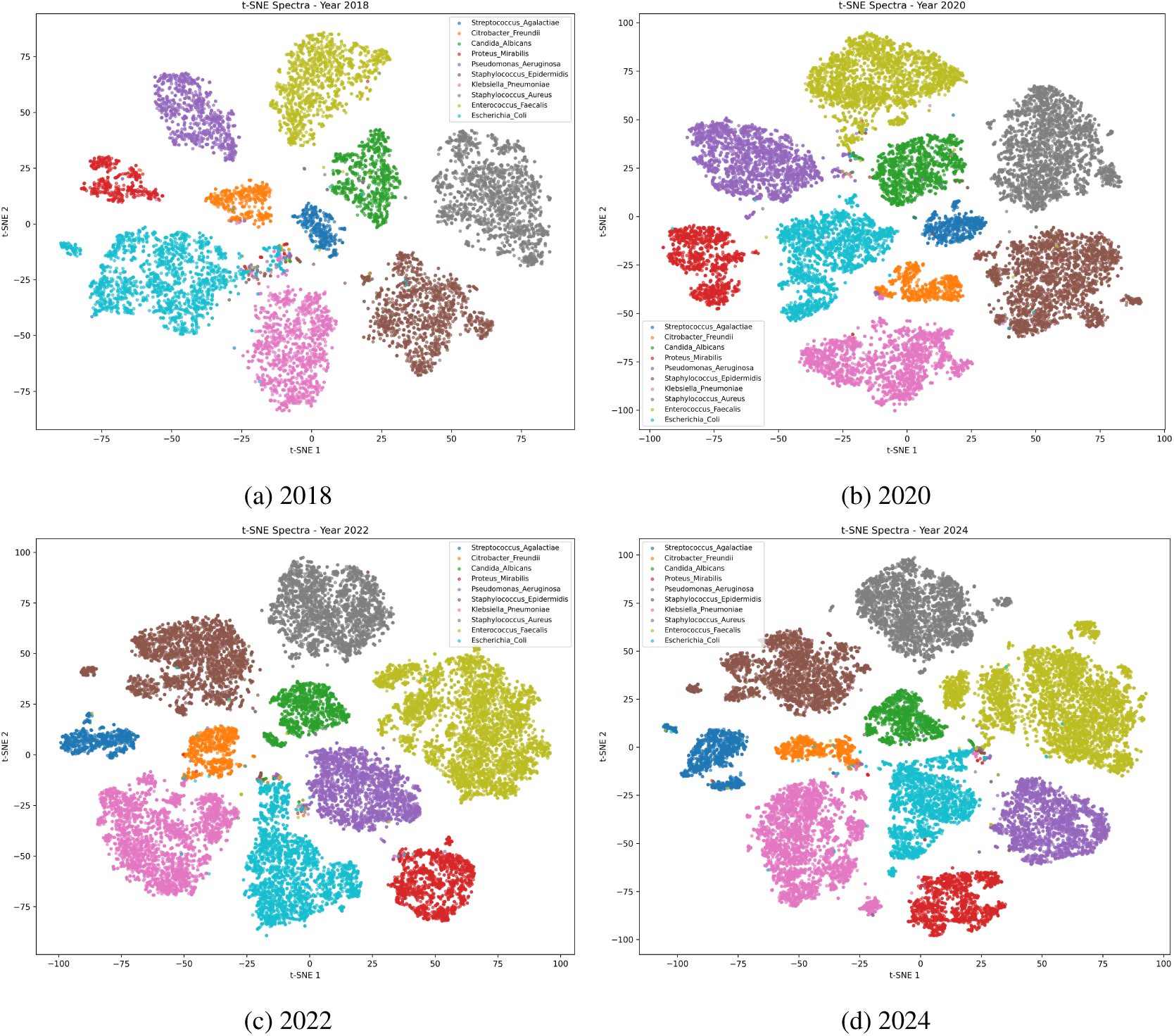
t-SNE embeddings of the ten most represented microorganisms in the dataset, plotted separately for some years. While the t-SNEs are computed independently and thus not aligned spatially, all years consistently exhibit ten separable clusters, reflecting robust inter-class separability across time.

Despite previous research conducted with partial subsets of this dataset, the complete dataset presented here has not yet been fully utilized or explored to its maximum potential. Similar to the impact of publicly available resources like the DRIAMS and RKI datasets [4, 7], we anticipate that this dataset will significantly advance the development and application of more sophisticated deep learning models, including Transformer-based approaches [10, 27], thereby fostering innovation in clinical microbiology research.

## Usage Notes

The *MARISMa* dataset is complemented with a set of Python scripts for data management and analysis. These scripts were designed to provide a practical introduction to working with the dataset and to facilitate initial experiments for bacterial identification within this dataset.

The script `simple_classifier.ipynb` offers a straightforward demonstration on processing and analyzing MALDI-TOF spectra, followed by a simple `Random Forest` classifier trained to discriminate between all available *Klebsiella* species within the selected time frame. This script is intended to inspire further analytical explorations.

## Data Availability

The *MARISMa* dataset supporting this work has been deposited in Zenodo and is publicly available at https://zenodo.org/records/17201597, with the version reviewed here being v2.0.0.

## Code Availability

All codes used for data preprocessing, cleaning, and preparation of the *MARISMa* dataset are publicly available in a dedicated GitHub^®^ repository (https://github.com/luciaschmidtsantiago/MARISMa). This repository also includes the scripts referenced in this work for classification tasks and other analyses.

## Acknowledgements

This work is partially supported by grant PID2023-146684NB-I00 funded by MCIN/AEI/10.13039/501100011033 and ERDF/EU and by the project TEC-2024/COM-89, by the project PI18/00997 from the Health Research Fund (FIS. Instituto de Salud Carlos III. Plan Nacional de I+D+I 2013-2016) of the Carlos III Health Institute (ISCIII, Madrid, Spain) partially financed by the European Regional Development Fund (FEDER) ‘A way of making Europe’. The funders had no role in the study design, data collection, analysis, decision to publish, or preparation/content of the manuscript. DRT (Sara Borrell CD22-00014) and BRS (Miguel Servet CPII19/00002) are funded by ISCIII. MBS is the recipients of the Intramural predoctoral contract 2022 from the Health Research Center of the Hospital Gregorio Marañón -IISGM-.

## Author Contributions

L.S.S., I.L.M., and J.M.M. processed, cleaned, and curated the dataset. D.R.T. and M.B.S. collected the clinical isolates from routine diagnostics. I.L.M. developed the software used for data analysis. D.R.T., B.R.S., and V.G.V. provided resources and acquired funding. A.G.L. wrote the initial manuscript draft and conducted code validation and addressed technical issues. All authors critically reviewed, edited, and approved the final version of the manuscript.

